# Refining Martini Force Field Interactions for Accurate Glycoprotein Modeling

**DOI:** 10.1101/2024.08.20.608764

**Authors:** Maziar Heidari, Mateusz Sikora, Gerhard Hummer

**Affiliations:** Department of Theoretical Biophysics, Max Planck Institute of Biophysics, Max-von-Laue Straße 3, 60438, Frankfurt am Main, Germany; Malopolska Centre of Biotechnology, Jagiellonian University, 30-387 Kraków, Poland; Institute of Biophysics, Goethe University Frankfurt, 60438 Frankfurt am Main, Germany

## Abstract

Covalently attached sugar molecules play important roles as mediators of biomolecular interactions. Molecular dynamics simulations are an indispensable tool to explore these interactions at the molecular level. The large time and length scales involved frequently necessitate the use of coarse-grained representations, which heavily depend on the parameterization of sugar-protein interactions. Here, we adjust the sugar-protein interactions in the widely used Martini 2.2 force field to reproduce the experimental second virial coefficients between sugars and proteins. In simulations of two model proteins in glucose solutions with adjusted force field parameters, we observe weak protein-sugar interaction. The sugar molecules are thus acting mainly as crowding agents, in agreement with experimental measurements. The procedure to fine-tune sugar-protein interactions is generally applicable and could prove useful also for atomistic force fields.

## Introduction

In biological systems, proteins interact closely with the many different osmolytes in the surrounding solution, ^1^ including amino acids, polyols and various sugars. As crowders, the osmolytes in the intracellular medium affect the protein-folding equilibria, protein stability, protein self-assembly and complex formation. ^2,3^ In cells, sugars are present not only in solution, but are also covalently attached to a majority of proteins through N-and O-glycosylation process.^4^ These protein modifications impact the intrinsic kinetics of protein interactions,^5^ protein dynamics^6^ and modulate the liquid-liquid phase separation (LLPS) of intrinsically disordered proteins (IDPs).^7^

Molecular simulations have been widely used to resolve the details of LLPS in atomistic and coarse-grained simulations. However, protein glycosylation is often ignored, yet glycans can have a substantial impact on the phase behavior and aging. ^8^ To accurately capture glycosylation effects, it is crucial to properly balance the energetics between sugars, solvents and proteins. Similarly, MD simulations have been used to explore the role of N-glycans in ligand binding,^9^ protein flexibility,^10^ and glycan shielding. ^11^ For the Martini force field, which is widely used for coarse-grained simulations of biological systems, ^12–15^ recent studies have shown that the protein-protein^16–18^ and osmolyte-osmolyte^19^ interactions have to be rescaled in order to correctly match the experimental observables. In simulations of complex biological systems, it is essential to accurately model cross interactions between components; however, the correct parameters for protein-sugar interactions remain largely unexplored. Here, we use Martini 2.2 and introduce a scaling factor to balance the cross interaction between proteins and glucose to match the experimentally determined values of the second viral coefficient for two model systems: cytochrome *c* and *α*-chymotrypsin dimers in glucose solution. With their “stickiness” properly adjusted, we then study the organization of glucose molecules around the proteins.

## Methods

We used the Martini 2.2 coarse-grained (CG) model^12^ to describe the energetics. We tuned the Lennard-Jones (LJ) interactions between sugar and proteins by adapting the procedure used previously to scale interactions between proteins^16,17^ and polysaccharides.^19^ Specifically, we rescaled the protein-sugar LJ interaction strengths *ϵ* with a *λ* parameter by setting *ϵ*_*λ*_ = *ϵ*_0_ + *λ*(*ϵ*_original_ − *ϵ*_0_). For *λ* = 1, the original cross interaction strength *ϵ*_original_ between sugars and proteins in the Martini model is recovered. For *λ* = 0, one obtains a very weak strength of *ϵ*_0_ = 2 kJ/mol corresponding to repulsive interactions. The scaling parameter *λ* is adjusted to reproduce the measured dependence of the second virial coefficient of a protein on the glucose concentration in the aqueous solution. We then validate *λ* by comparing the predicted effect of glucose on the dimerization equilibrium for a different protein to experiments.

We computed the second virial coefficients from the cumulative radial distribution function (RDF) between two components (glucose and protein). The McMillan and Mayer expression^20^

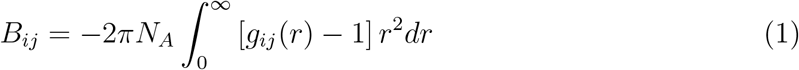

relates the second virial coefficient *B*_*ij*_ for solutes *i* and *j* to their RDF *g*_*ij*_(*r*) integrated over the distance *r*, with *N*_*A*_ being the Avogadro constant. Formally, the integral in eq 1, converges in the limit of large systems and in a grand canonical ensemble. ^21^ For finite systems, however, radial distribution functions *g*_*ij*_(*r*) can noticeably deviate from one,^21,22^ which will lead to a cubically growing divergence of *B*_*ij*_. To address these challenges, we used systems of comparably large size and compared the results obtained for different sizes. In addition, we explicitly corrected for any remaining small deviations of the RDF from one at large distances. For this, we averaged the RDF tail over a distance window of width Δ,

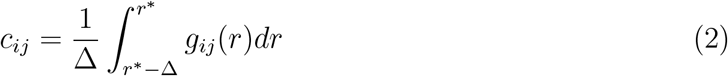

with *r*\* being a distance larger than the correlation length. We then replaced the ideal RDF large-distance limit of one in eq 1 by this average to estimate the second virial coefficient *B*_*ij*_ as

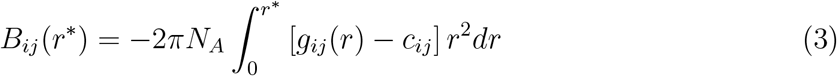

We compared results for *B*_*ij*_(*r*\*) obtained for large integration cutoffs *r*\* to experimental values of protein-glucose cross second virial coefficients (*B*_23_) and glucose-dependent dimerization constants obtained by Pielak and coworkers. ^23,24^

### Dimer dissociation

We estimate the rate of *α*-chymotrypsin dimer dissociation from the statistics of dissociation events in multiple simulation runs starting from a dimerized state. For this, we use a maximum-likelihood estimator,

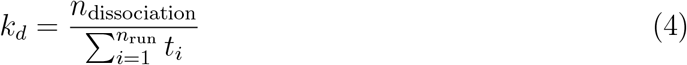

where *n*_dissociation_ ≤ *n*_run_ is the number of dissociation events observed in the *n*_run_ runs and *t*_*i*_ is the time point of the event or, if no dissociation occurred, the duration of the run. The denominator is thus the aggregate time in the bound state across the different runs.

## Simulation Models

As model systems, we used the structures of *Saccharomyces cerevisiae* iso-1 cytochrome *c* (PDB ID: 2ORL;^25^ non-heme ligands removed) and *Bos taurus α*-chymotrypsin dimer (PDB ID: 4CHA^26^). We used the martinize.py script^13,27^ to convert the protein structures into coarse-grained representation within the Martini 2.2 scheme. ^12^ We used the DSSP algorithm to assign the secondary structure restraints. ^28^ We used Elastic Network in Dynamics (ElNeDyn) to generate elastic bonds within the ordered domains to maintain the tertiary structure, ^29^ with default parameters settings, a distance range of 0.9 nm and a force constant of 500 kJ*/*(mol nm^2^). All systems were solvated with coarse-grained water containing 10% anti-freeze (WF) particles.^12^ Charge neutrality and salt molarity were established by replacing water particles with ions following standard GROMACS procedure. We used Cl^−^ ions to neutralize the cytochrome *c* systems so as to simulate the experimental deionized water condition.^23^ We added Na^+^ and Cl^−^ ions to match the experimental concentration of 200 mM NaCl^24^ in overall neutral *α*-chymotrypsin systems. We used the GROMACS 2020.6 software package to simulate all systems^30^ and prepared visualizations using the Visual Molecular Dynamics (VMD) software.^31^

Cytochrome *c* binds heme C within a dedicated pocket. With heme B parameters available in the Martini force-field, we used heme B instead of heme C to construct the holo state of the protein.^32^ In heme C, the two vinyl groups of heme B are replaced by thioether linkages, which covalently attach the heme to two cysteine residues of cytochrome *c*. Being largely buried within the near-rigid protein scaffold, the differences between the two hemes are negligible for the purpose of our calculations. To mimic the low experimental pH of 3.5 of the *α*-chymotrypsin dimer solution, ^24^ we used PROPKA^33,34^ with the all-atom AMBER ff99 energy function^35^ and the CHARMM-GUI server^36,37^ to determine amino acid protonation states. We found that six amino acids were protonated in each monomer (see Figure S1). We modified the protonated residues by adjusting their charge accordingly and changing the type of the side chain particle in the Martini force field, i.e., from Qa to P3 for ASP, from Qa to P1 for GLU and from P1 to Qd for HIS.

We performed energy minimization on the initial models using the steepest descent algorithm with an energy minimization step of 0.01 nm and a force tolerance of 1000 kJ mol^−1^ nm^−1^. This was followed by 150 ns of MD simulation with a time step of *δt* = 0.015 ps at a tem-perature of 300 K established using a velocity rescaling thermostat ^38^ with a time constant of 1 ps. Then we performed 750 ns MD simulation in an isothermal-isobaric ensemble at a temperature of 300 K established using a velocity rescaling thermostat and a pressure of 1 bar established using an isotropic Berendsen barostat^39^ with time constant 12 ps and compressibility 3 *×* 10^−4^ bar^−1^. For production runs, we used a velocity rescaling thermostat and an isotropic Parrinello-Rahman barostat^40^ with identical time constants. The time step of production runs was set to *δt* = 0.03 ps. The production runs were at least 60 *μ*s for glucose in solution and 30 *μ*s for all other cases. In all simulations we used cubic boxes of size *L* ≈ 30 nm or *L* ≈ 40 nm. The concentrations of glucose for cytochrome *c* and *α*-chymotrypsin systems were set to 0.5 M (cytochrome *c*; Figure 1a) and 0.1 M (*α*-chymotrypsin monomer and dimer; Figure 2a), respectively. The RDFs were computed between the center of mass of protein and sugar molecules using the GROMACS built-in command *gmx rdf*. The bin size, maximum range and configurational sampling frequency of RDFs were set to 0.1 nm, 10 nm and 1.5 ns, respectively. The RDFs were orientationally averaged about the proteins. Previous studies have introduced optimum Lennard-Jones scaling parameters *α* = 0.3 (ref 16) and *γ*=0.5 (ref 19) for protein-protein and sugar-sugar interactions, respectively. Here, we kept the prescribed sugar-sugar interaction scaling, but consistently used *α* = 0.7 scaling for the optimal protein-protein interaction as we determined before^17^ (see Figures S2 and S3 for validation of the earlier findings). However, we varied the interaction scaling factor *λ* of protein-sugar interactions. For the RDF tail correction of sugar-sugar interactions, we used Δ = 1 nm and *r*\* = 5 nm; and for the sugar interactions with the proteins cytochrome *c* and *α*-chymotrypsin, we used Δ = 2 nm and *r*\* = 10 nm.

**Figure 1:**
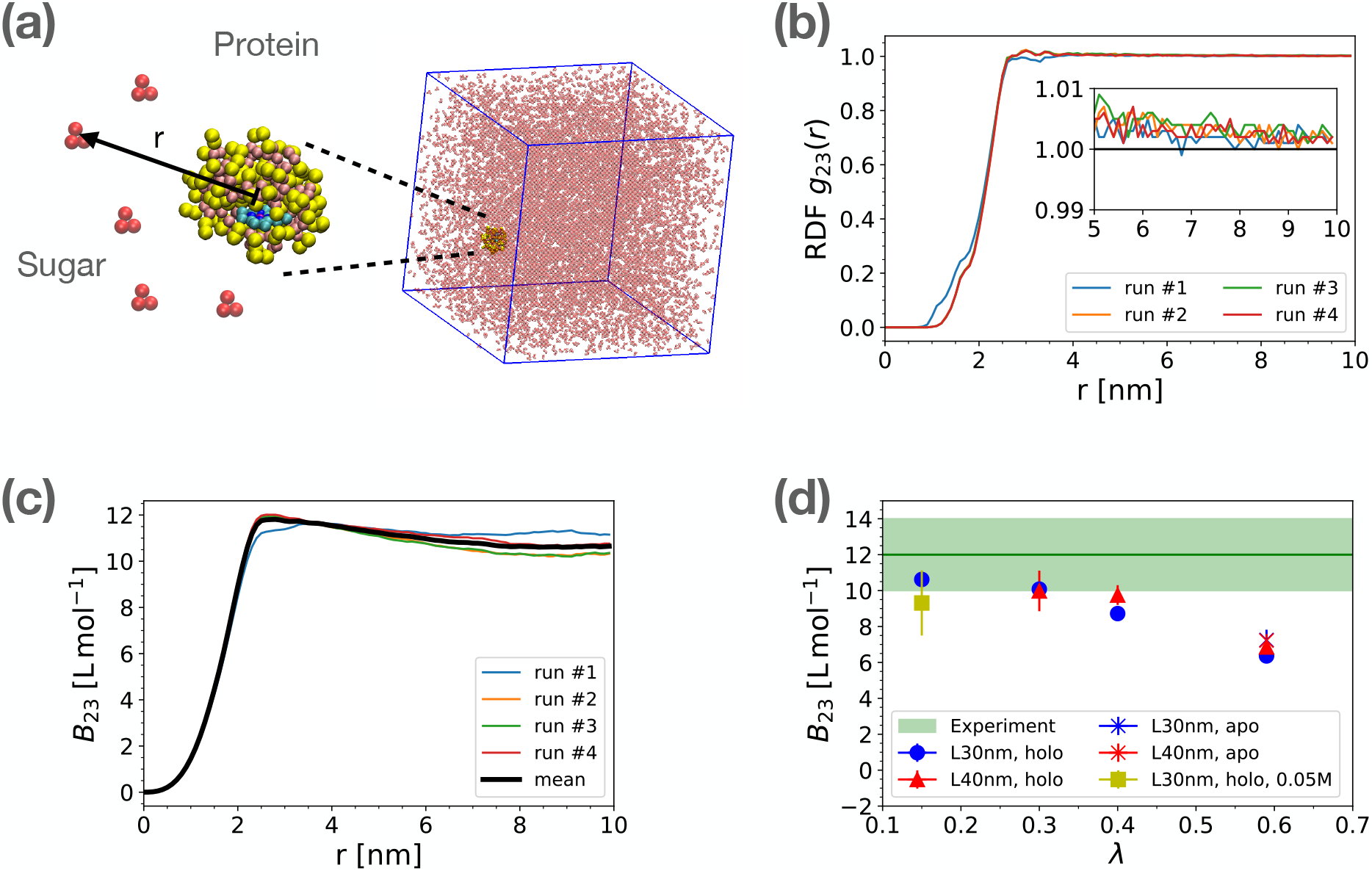
Balancing protein and sugar interactions. (a) Coarse-grained model of glucose (red beads) and cytochrome *c* in holo state (yellow and pink beads). The heme C molecule is shown in the pocket of cytochrome (cyan). The simulation box size is 30 nm and the glucose concentration is 0.5 M. (b) Radial distribution function of glucose molecules about the cytochrome *c* protein for scale factors *γ* = 0.5 and *λ* = 0.15. The distance is measured between the center of mass of the two components. The results of four independent runs are shown with different colors. The inset zooms in on the convergence region of RDFs within the range of 5 nm to 10 nm. (c) Second virial coefficient of protein-sugar interaction calculated by integrating RDFs in panel b according to eq 3 for *λ* = 0.15. (d) Second virial coefficient of cytochrome *c* and glucose as a function of cross interaction scaling parameter *λ*. The green shaded area shows the experimental value ^23^ with standard error. The symbols in panel d show averages of *B*_23_(*r*\*) over the range 8 nm *< r*\* *<* 10 nm and across the different runs, with error bars indicating standard errors of the mean. For *λ* = 0.15, also a system with 0.05 M glucose was studied (filled square) instead of the 0.5 M in the other systems. Results are shown for holo (filled symbols) and apo cytochrome *c* (stars) simulated in boxes of size *L* ≈ 30 nm (blue) and 40 nm (red).

**Figure 2:**
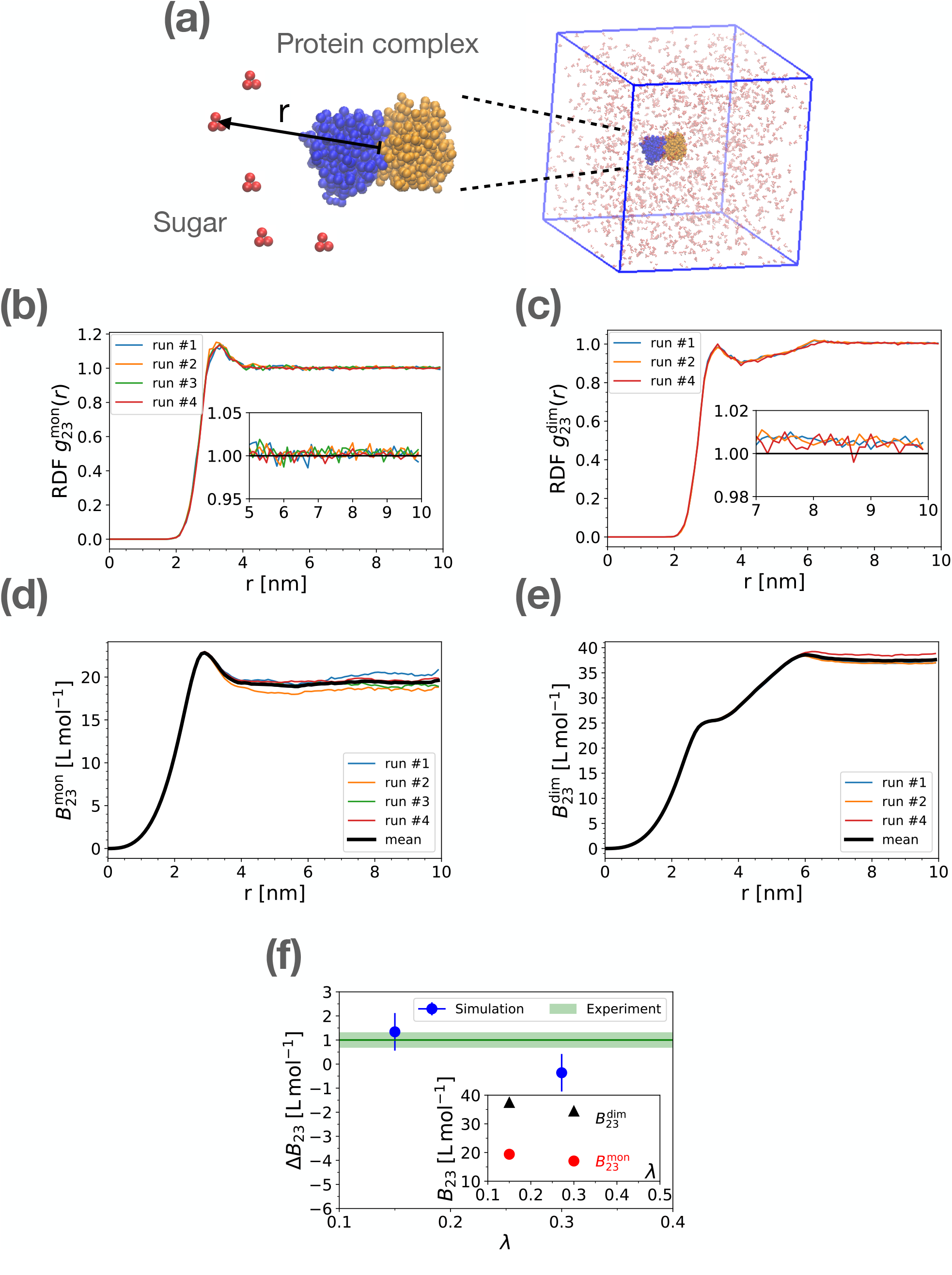
Balancing protein and sugar interactions. (a) Coarse-grained model of glucose (red beads) and *α*-chymotrypsin dimer (PDB ID: 4CHA) with monomers colored in blue and orange. The simulation box size is 30 nm and the glucose concentration is 0.1 M. (b, c) Radial distribution function of glucose molecules about *α*-chymotrypsin monomer and dimer with scale factors *α* = 0.7, *γ* = 0.5 and *λ* = 0.15 for protein-protein, sugar-sugar and protein-sugar interactions. The distance is measured between the centers of mass of the components. The results of independent runs are shown with different colors. Insets zoom in on the convergence region of RDFs within the range of 5 nm to 10 nm. (d, e) Convergence of second virial coefficient of sugar and protein in monomer 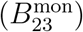 and dimer 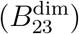 configurations using integration of RDFs of panels b and c according to eq 3. (f) Effect of glucose on the stability of *α*-chymotrypsin dimer, 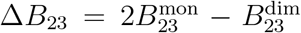, as a function of protein-sugar interaction scaling parameter *λ*. The green shaded area shows the experimental value ±SE, which matches with the simulation results at *λ* = 0.15. The inset shows the second virial coefficients of sugar and *α*-chymotrypsin in monomer and dimer states. In dimer *α*-chymotrypsin, the result of simulation run #3 is not shown as the dimer dissociated near the beginning of the run (see Figure S8a). The symbols and error bars in panel f are the average and standard error of means obtained by averaging Δ*B*_23_(*r*) over the range 8 nm *< r <* 10 nm for each run.

## Results

### Second osmotic virial coefficient for glucose-cytochrome *c* interaction

To determine the preference of the glucose molecules to bind to cytochrome *c*, we first analysed the RDFs of glucose around cytochrome *c*. In Figure 1b we show examples of four independent simulation runs at *λ* = 0.15. We found that the excluded radius extends approximately 1 nm from the protein center. Between 3 and 4 nm, close to the protein surface, the glucose molecules form a very weakly structured layer. Beyond, the RDF approaches one, the ideal gas limit. The inset zooms in on the convergence of RDFs, highlighting the small but noticeable deviations from one, as estimated by *c*_*ij*_ in eq 2. Similar RDFs are found for *λ* = 0.3 (Figure S4a,b), with the structured layers enhanced for larger *λ* values (Figure S4c-f). The cross osmotic second virial coefficient *B*_23_ for glucose and protein calculated from different simulation runs converge at distances of *r*\* ≈ 8 to 10 nm (Figure 1c). To find the optimal value of *λ* corresponding to the experimentally measured values of *B*_23_, we looked into the dependence of *B*_23_ on *λ* (Figure 1d). A monotonic decrease of *B*_23_ with increasing *λ* is consistent with the rise in attractive interactions between protein and sugar. The calculated *B*_23_(*λ* = 0.15) = 10.63 ± 0.33 L mol^−1^ matches the experimental value of 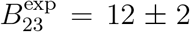 L mol^−1^,^23^ independent of the simulation box size (compare filled circles and triangles in Figure 1d). To test how the results depend on the presence of the heme ligand, we repeated simulations with heme B removed from the cytochrome *c* pocket (apo state). This led to a very small change in *B*_23_ (star symbols in Figure 1d) compared with the holo state. As shown in Figure S6, in the apo state glucose molecules can partially penetrate the binding pocket, leading to a higher first peak in the RDF. As a test of a possible glucose concentration dependence, the *B*_23_ value obtained in simulations with a reduced glucose concentration of M is statistically consistent with the results obtained for 0.5 M glucose (Figures 1d and S7). In summary, after scaling the glucose-protein interactions with a factor of *λ* = 0.15, we obtained a *B*_23_ value in agreement with measurements on cytochrome *c*. For higher values, *λ* ≥ 0.4, the calculated *B*_23_ is noticeably too small.

### Glucose effect on *α*-chymotrypsin dimerization

To validate the scale factor *λ* = 0.15, we turned to our second model system, *α*-chymotrypsin. For this protein, the effect of added glucose on the monomer-dimer equilibrium has been characterized by analytical ultracentrifugation.^24^ The apparent dissociation constant *K*_*d*_(*C*) depends on sucrose concentrations *C* as 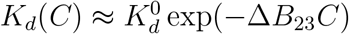 with 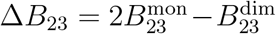 the difference in glucose-protein second virial coefficients for two protein monomers and a dimer. The experiments found a stabilizing effect for glucose ^24^ with Δ*B*_23_ ≈ 1.0 0.3 M^−1^. To calculate this effect for our Martini model with scaled sugar-protein interactions, we first computed the RDF of glucose and the protein in monomer 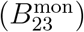 and dimer 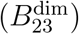 states separately, as shown in Figure 2b,c. During the simulations, we observed several events of spontaneous dissociation of *α*-chymotrypsin dimers (see Figure S8). Thus, for the computation of RDFs in the dimer state, we considered only the initial time interval during which the monomers were associated. In contrast to the RDFs of cytochrome *c* at *λ* = 0.15, we found that glucose molecules are more attracted to the *α*-chymotrypsin monomer (see more prominent peak at *r* = 3 nm in Figure 2b. The decrease in the RDF of glucose around the dimer complex close to *r* = 4 nm can be explained by the asphericity of the dimer (Figure 2c). The RDFs of monomer and dimer at *λ* = 0.3 also show similar trends (Figure S9). The effect of glucose on the stability of the *α*-chymotrypsin dimer can be quantified and compared with experimental values ^24^ using 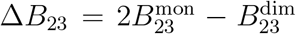. We find reasonable convergence of *B*_23_ at *r* above 5 and 7 nm for monomer and dimer, respectively (see Figure 2d,e). Similar to cytochrome *c, B*_23_ decreases with *λ* for both monomer and dimer (see inset in Figure 2f). Reassuringly, Δ*B*_23_ for *λ* = 0.15 matches the experimental value for *α*-chymotrypsin (Figure 2f).

Having obtained an optimal scaling parameter *λ* = 0.15 for the sugar-protein interactions in Martini 2.2, we next set out to determine how the relatively weak sugar-protein interactions govern the arrangement of glucose molecules in the vicinity of the two proteins. We calculated the cumulative distribution of the average number of contacts between glucose molecules and the residues of cytochrome *c* as shown in Figure 3. Consistent with the described *B*_23_ dependence on *λ*, we saw only very transient contacts that significantly decrease with reducing *λ* values.

**Figure 3:**
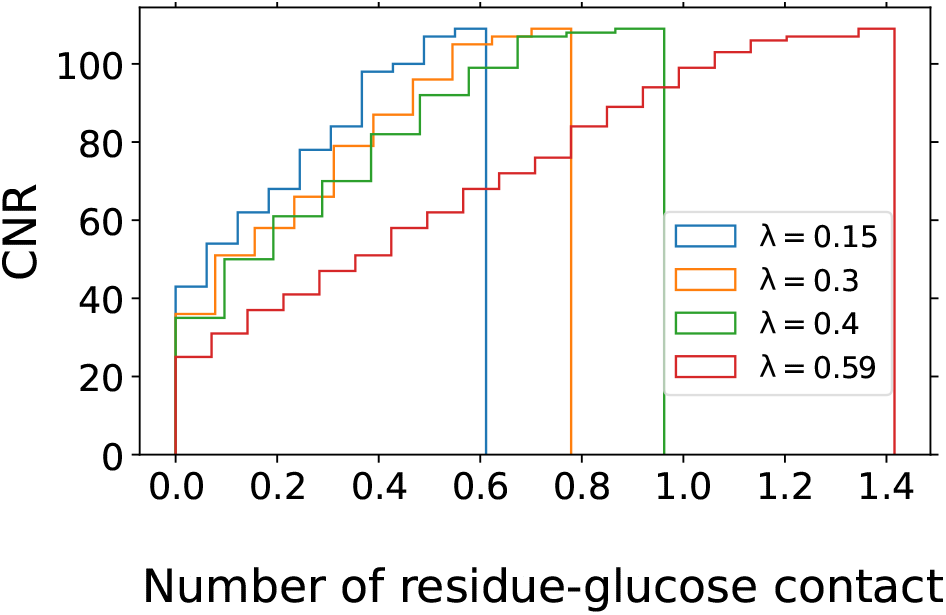
Glucose-protein contact number. Shown is the cumulative number of residues (CNR) in holo cytochrome *c* as a function of their mean number of contacting glucose beads, i.e., the number of amino acids that have less than the number of residue-glucose contacts given on the *x* axis. A glucose molecule is considered to be in contact with a residue if any glucose bead is within 0.94 nm of the residue center of mass. The steps at zero reflect the fact that for *λ* = 0.15, 0.3, 0.4 and 0.59, there are 6, 11, 15 and 19 residues without any glucose contacts during the simulations, respectively. The CNRs were obtained using the average number of contacts across four independent simulation runs.

### Dimer dissociation

In our MD simulations, we observed two *α*-chymotrypsin dimer dissociation events at times *t*_1_ = 3 *μ*s and *t*_2_ = 17 *μ*s (Figure S7a). In the other two runs, the dimer stayed bound for the duration of the simulations, *t*_3_ = 46 *μ*s and *t*_4_ = 45 *μ*s.

The maximum-likelihood estimate for the dissociation rate, eq 4, is *k*_*d*_ ≈ 1*/*(55.5 *μ*s). The measured value^24^ of the dimer dissociation constant is approximately *K*_*d*_ ≈ 1*/K*_2,app_ ≈ 20 *μ*M. If we assume diffusion-limited association, this value of *K*_*d*_ would correspond to an experimental dissociation rate of approximately *k*_*d*_ ≈ 10^9^ M^−1^ s^−1^ *K*_*d*_ ≈ 1*/*(50 *μ*s). Clearly, this near-perfect agreement is somewhat fortuitous, considering the approximations involved and the use of a coarse-grained simulation model. Nevertheless, it is reassuring that also the protein-protein interaction comes out about right for the scale factor *α* = 0.7 used here.

## Conclusion

We developed an approach to accurately model sugar-protein interactions in the Martini coarse-grained scheme by scaling the cross interaction strength between sugar and protein beads. We determined an optimal scale factor *λ* = 0.15 by matching the calculated and measured^23^ second virial coefficients for glucose and cytochrome *c*. With this parameter, we then recovered the effect of glucose on the dimerization equilibrium of *α*-chymotrypsin.^24^

The small value of the scaling parameter implies a very weak interaction between the sugar and protein, which is in-line with the general picture that the sugar molecules act as inert crowders and interact only weakly with proteins. We expect this effect to be important also for larger saccharides and glycans, as the osmotic coefficient is expected to grow with the degree of polymerization.^19,41^ The weak attractions between sugars and proteins contribute to the inhibitory effect of glycans on protein-protein interactions, e.g., to create a ‘glycan shield’ around antigens that protect against antibody binding. ^42^

As a general issue, we conclude that simple mixing rules such as Lorentz-Berthelot, should not be expected to give accurate renderings of binding and phase behavior. Even if AA and BB interactions between two compounds A and B are well balanced (here, proteins and sugars in aqueous solution), there is no guarantee that a predefined mixing scheme will be sufficiently accurate for the AB interactions. We showed that by simply using the geometric average of protein-protein (*α* ≈ 0.7; ref 17) and sugar-sugar scale factors (*γ* = 0.5; ref 19) for protein-sugar interactions, 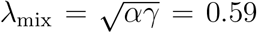, the sugar molecules are too sticky and adhere to the protein surface. Even with a lower value of *α* = 0.3 for protein-protein interactions,^16,19^ the sugar-protein scale factor expected from mixing, *λ* ≈ 0.39, is larger than our optimal value of *λ* = 0.15.

Here, we focused on the Martini 2.2 scheme; however, we expect similar reasoning can be made for Martini 3. As the Martini force field has paved the way for simulations of large length-scale and long time-scale problems, we believe that our scaling of sugar-protein interaction is a critical step on the path to whole-cell simulations^43^ as many cellular components interact either with the solution glycans or with post-translationally glycosylated components.

## Acknowledgments

The authors thank Sören von Bülow for providing the python script that rescales the cross interaction strength between different bead types in the Martini 2.2 force field. This work was supported by the Max Planck Society and by CRC 1507 ‘Membrane-associated protein assemblies, machineries and supercomplexes’ of the Deutsche Forschungsgemeinschaft (DFG project number 450648163). We thank the Max Planck Computing and Data Facility (MPCDF) for computational support. M.S. receives support through the Dioscuri program initiated by the Max Planck Society, jointly managed with the National Science Centre in Poland, and mutually funded by the Polish Ministry of Education and Science and German Federal Ministry of Education and Research (UMO-2021/03/H/NZ1/00003)

## Supporting Information Available

Supporting Figures S1 to S9.

## Supporting Information

### Supporting Figures

**Figure S1:**
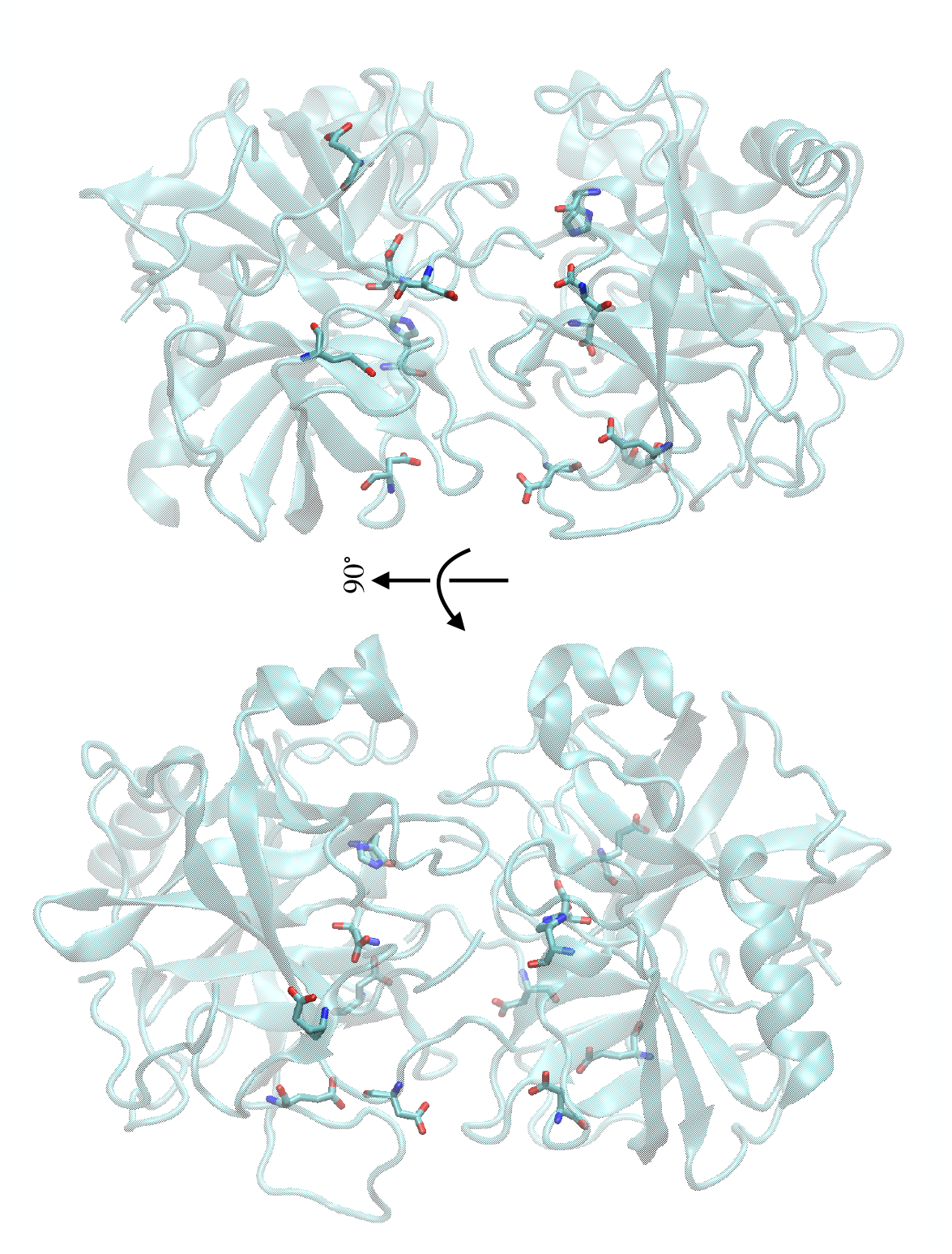
Protonated residues of *α*-chymotrypsin at pH 3.9 identified using PROPKA^S1,S2^ with all-atom AMBER ff99^S3^ and CHARMM-GUI server. ^S4,S5^ In each monomer, GLU20 (pKa = 5.48), HIS57 (pKa = 8.37), ASP64 (pKa = 4.67), GLU70 (pKa = 6.91), ASP153 (pKa = 5.16), and ASP194 (pKa = 3.98) were protonated. The protonated residues are shown as liquorice, and the protein dimer is shown as cartoon. The protonated residues are concentrated at the interface between the monomers.

**Figure S2:**
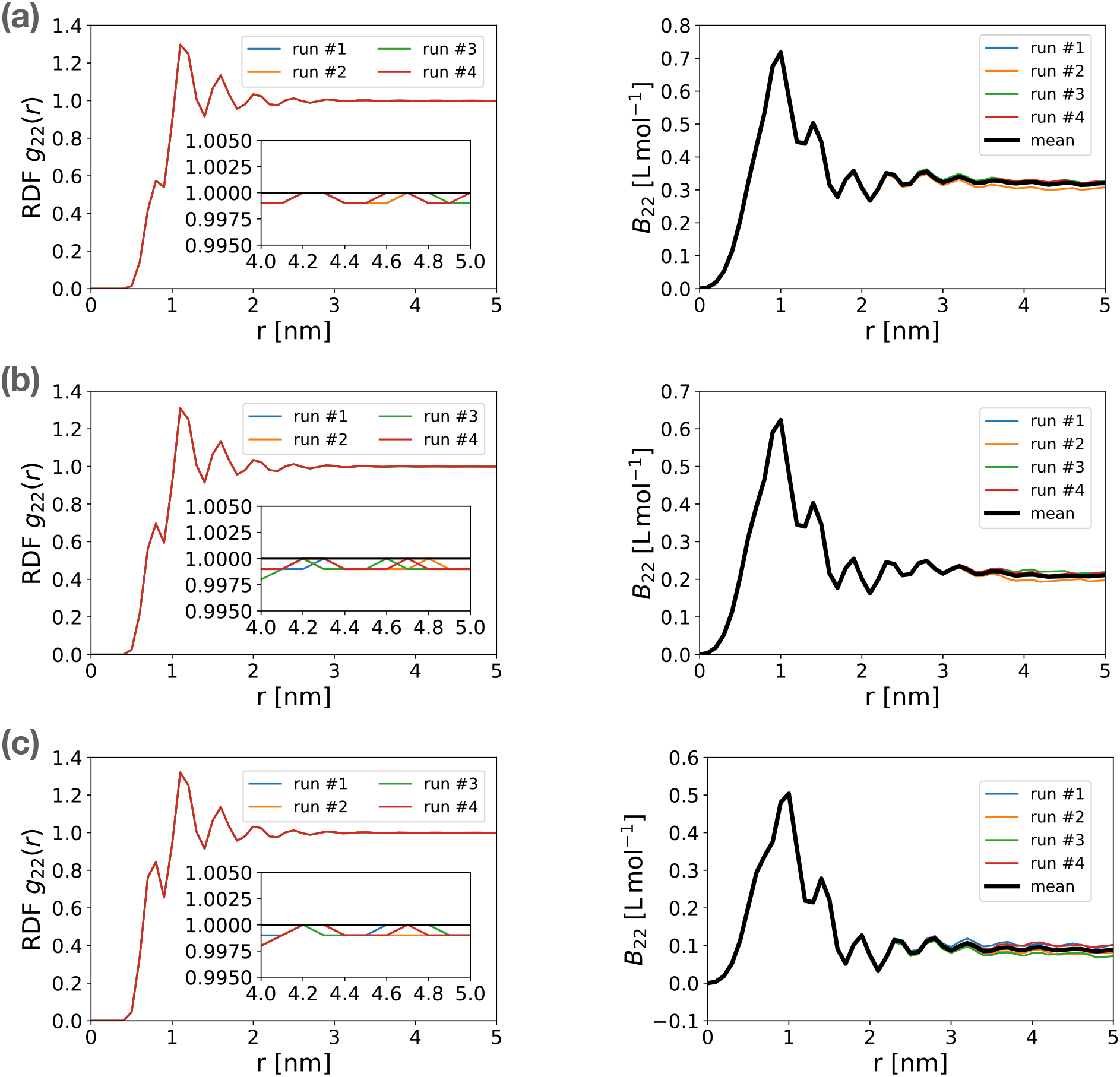
Glucose-glucose radial distribution function (RDF) and convergence of osmotic second virial coefficient. RDFs were computed between the respective centers of mass. In each panel, the RDF (left) and *B*_22_ (right) of four independent runs are shown. The scaling parameter between sugar-sugar interaction (*γ*) is *γ* = 0.4 (a), *γ* = 0.5 (b) and *γ* = 0.6 (c). The concentration of the sugar molecules is 0.1 M and the system is a cube of size *L* ≈ 30 nm. The inset zooms in on the RDF tails. To compute *B*_22_, we used Δ = 1 nm and *r*\* = 5 nm.

**Figure S3:**
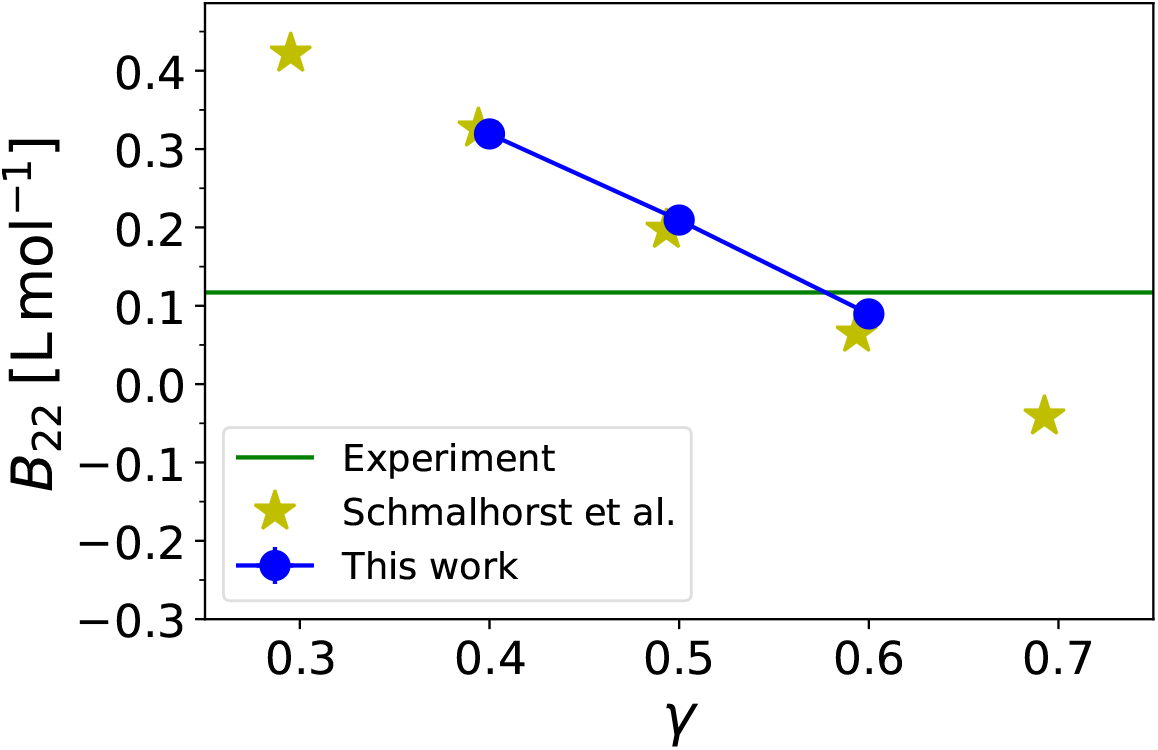
Osmotic second virial coefficient of glucose-glucose interaction. *B*_22_ is shown against scaling parameter *γ* using RDFs at concentrations 0.1M of glucose solutions. The results of Schmalhorst et al. ^S6^ are shown by yellow stars.

**Figure S4:**
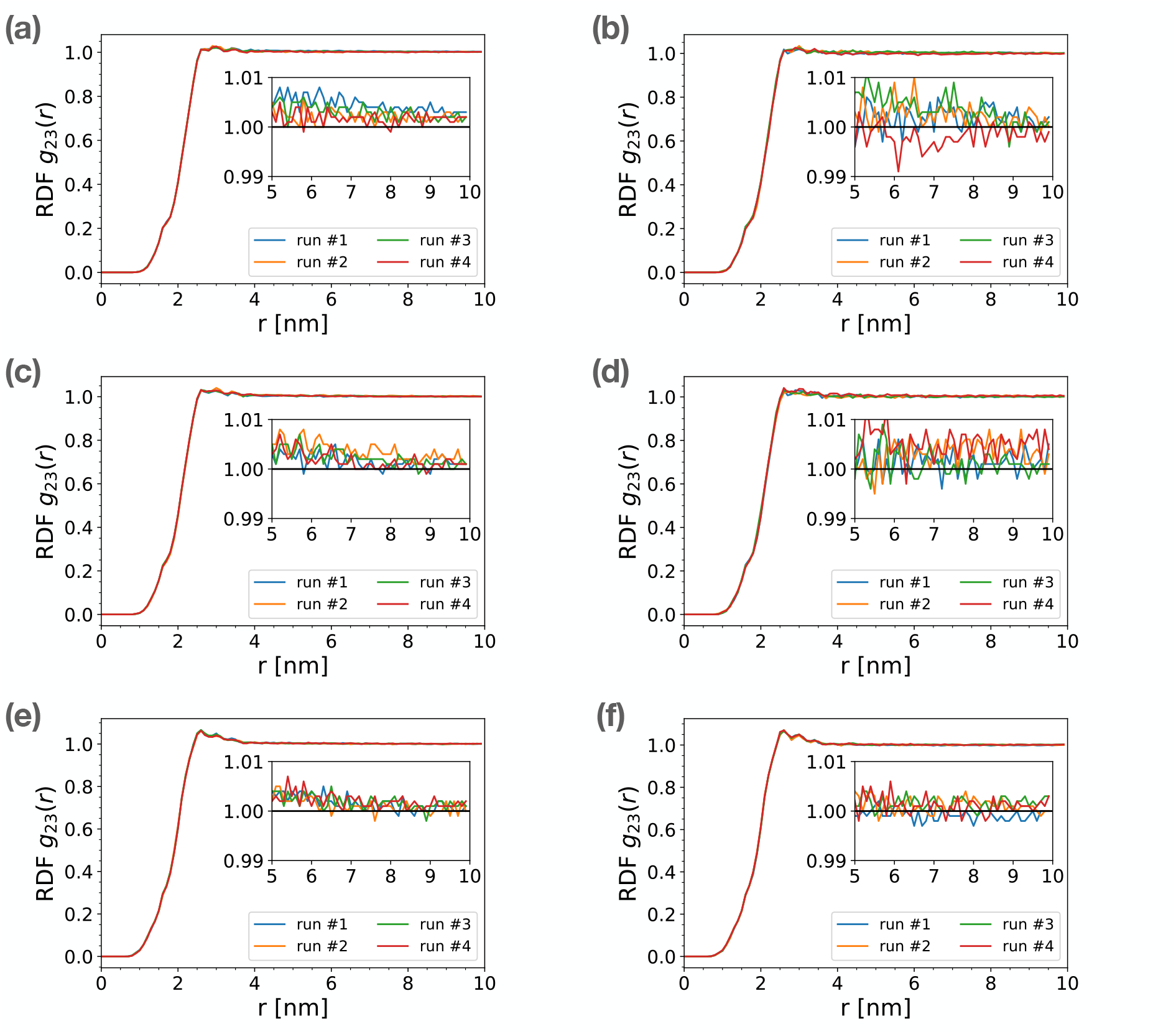
Radial distribution function (RDF) of glucose around holo cytochrome *c* at a glucose concentration of 0.5 M. The RDFs were computed using different protein-sugar scaling parameters (*λ*) and simulation box sizes (*L*). (a, b) *λ* = 0.3; (c, d) *λ* = 0.4; (e, f) *λ* = 0.59; (a, c, e) *L* = 30 nm; (b, d, f) *L* = 40 nm. Insets zoom in on the RDF tail in the range of 5 to 10 nm.

**Figure S5:**
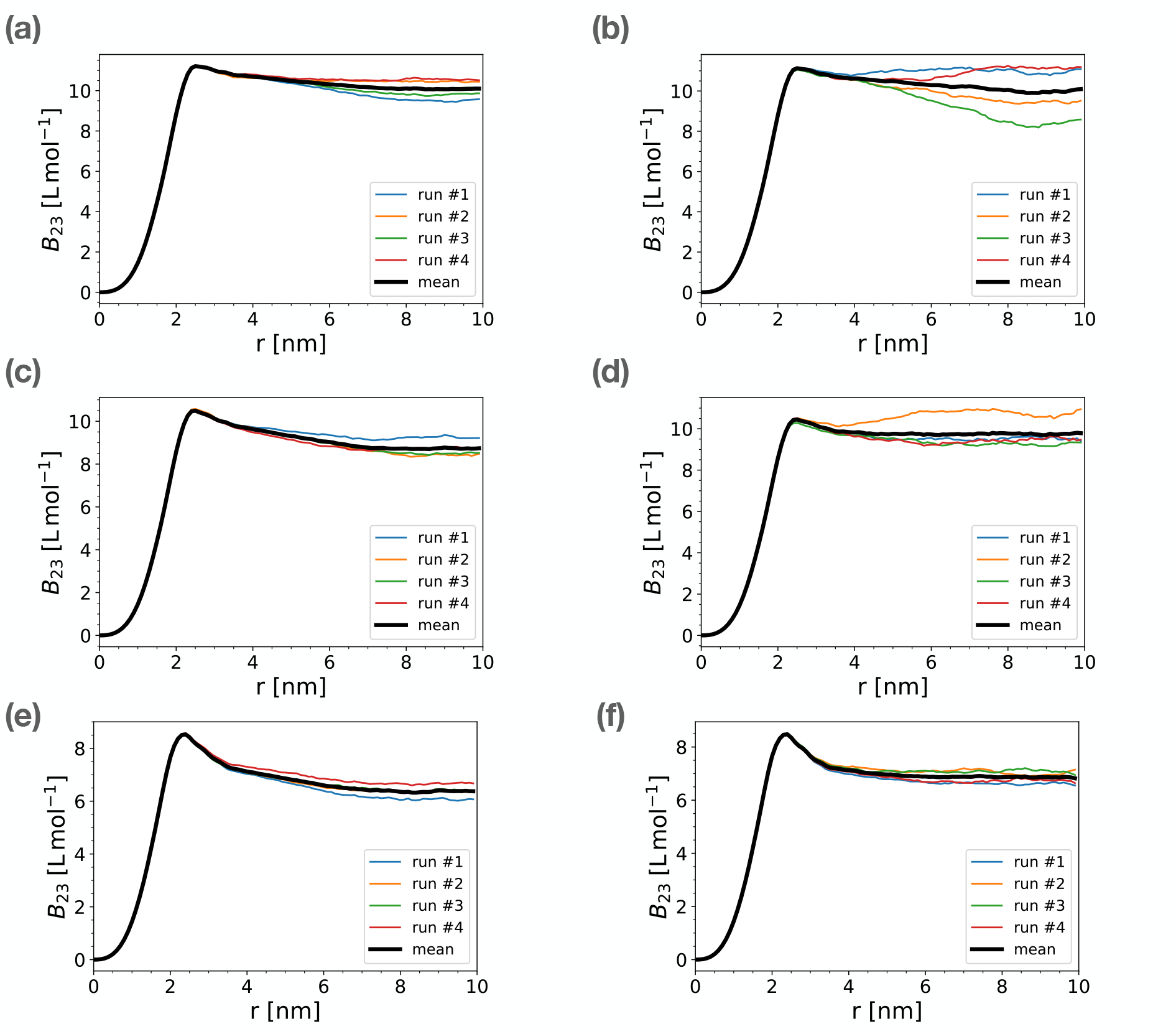
Convergence of osmotic second virial coefficient (*B*_23_) for the interaction between glucose and holo cytochrome *c* at a glucose concentration of 0.5 M. The *B*_23_ were computed using different protein-sugar scaling parameters (*λ*) and simulation box sizes (*L*). (a, b) *λ* = 0.3; (c, d) *λ* = 0.4; (e, f) *λ* = 0.59; (a, c, e) *L* = 30 nm; (b, d, f) *L* = 40 nm. For each panel, the results of four independent simulation runs are included and the thick black curve shows the average of four runs. To compute *B*_23_, we used Δ = 2 nm and *r*\* = 10 nm.

**Figure S6:**
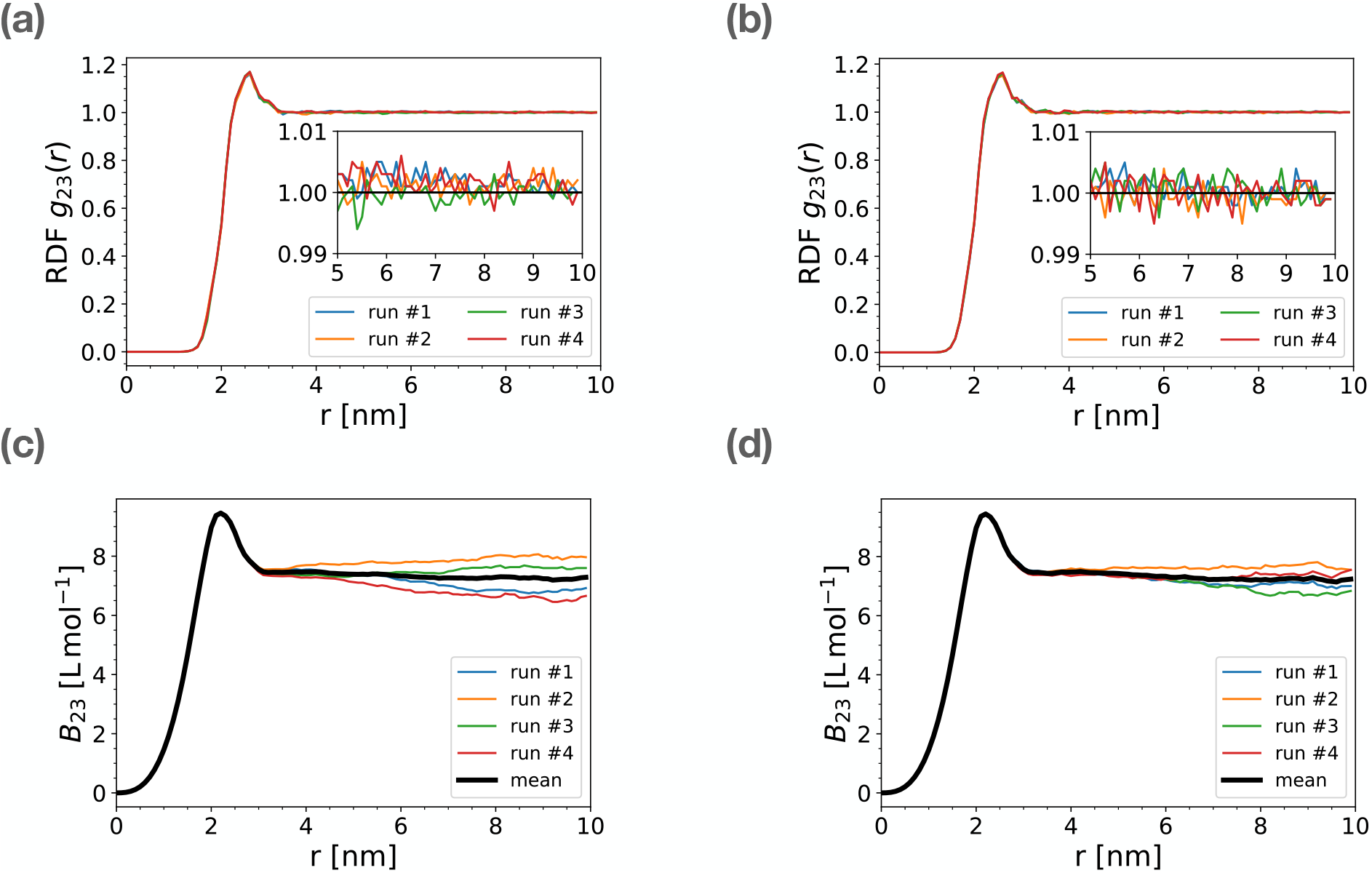
Radial distribution function (RDF) of glucose around apo cytochrome *c* at a glucose concentration of 0.5 M, and convergence of the associated osmotic second virial coefficient (*B*_23_). The RDFs and *B*_23_ were computed using protein-sugar scaling parameter (*λ* = 0.59) and simulation box size *L* = 30 nm (a, c) and *L* = 40 nm (b, d). Insets zoom in on RDF convergence in the range 5 to 10 nm. To compute *B*_23_, we used Δ = 2 nm and *r*\* = 10 nm.

**Figure S7:**
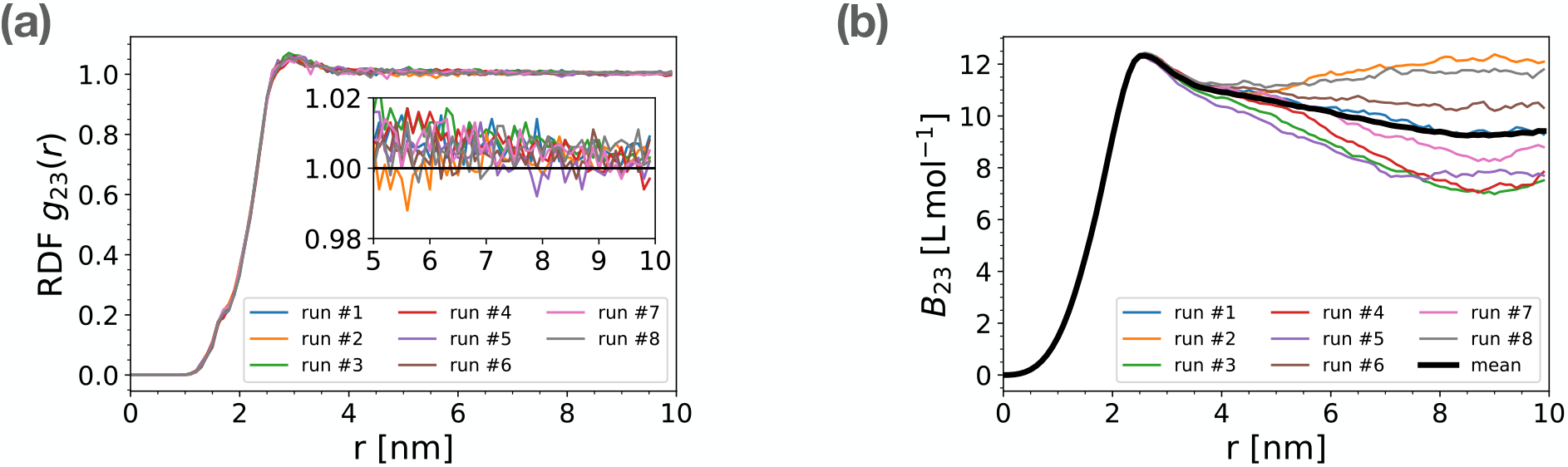
Radial distribution function (RDF) of glucose at a glucose concentration of 0.05 M around holo cytochrome *c*, and convergence of the associated osmotic second virial coefficient (*B*_23_). The RDFs (a) and *B*_23_ (b) were computed using a protein-sugar scaling parameter *λ* = 0.15 and a simulation box size of *L* = 30 nm. Insets zoom in on RDF convergence in the range 5 to 10 nm. To compute *B*_23_, we used Δ = 2 nm and *r*\* = 10 nm.

**Figure S8:**
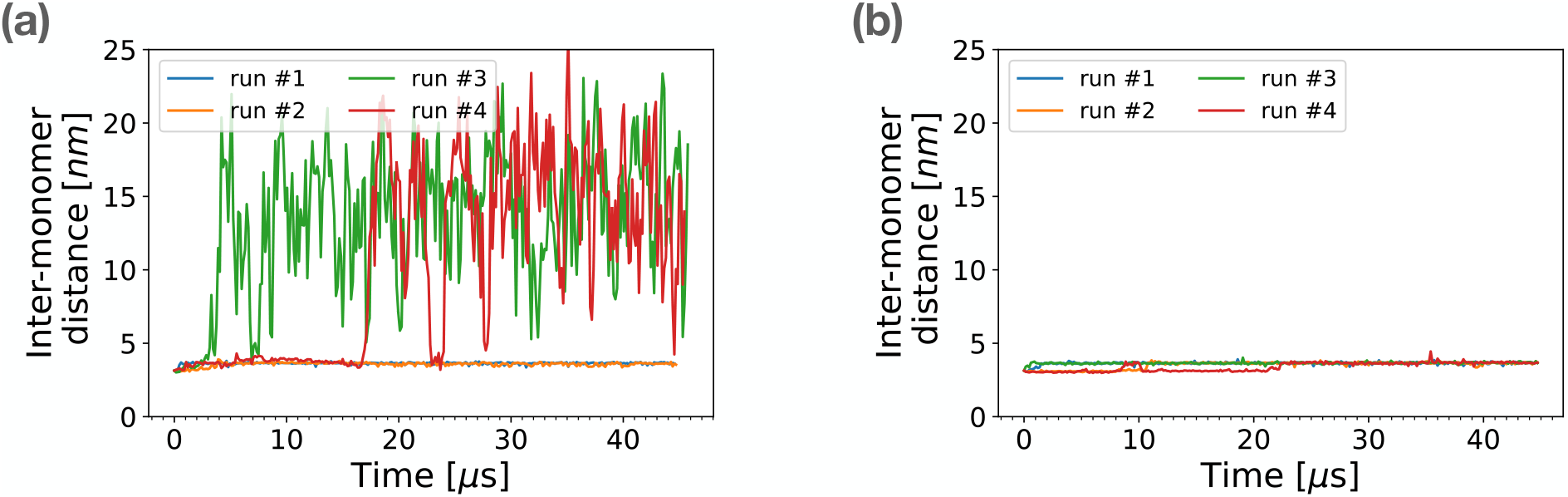
Inter-monomer distance of two *α*-chymotrypsin proteins starting from a dimeric state. The time series of the distance between monomer centers of geometry in dimer *α*-chymotrypsin is plotted for *λ* = 0.15 (a) and *λ* = 0.3 (b). Protein-protein interactions were scaled with *α* = 0.7 (ref S7). The glucose concentration was 0.1 M.

**Figure S9:**
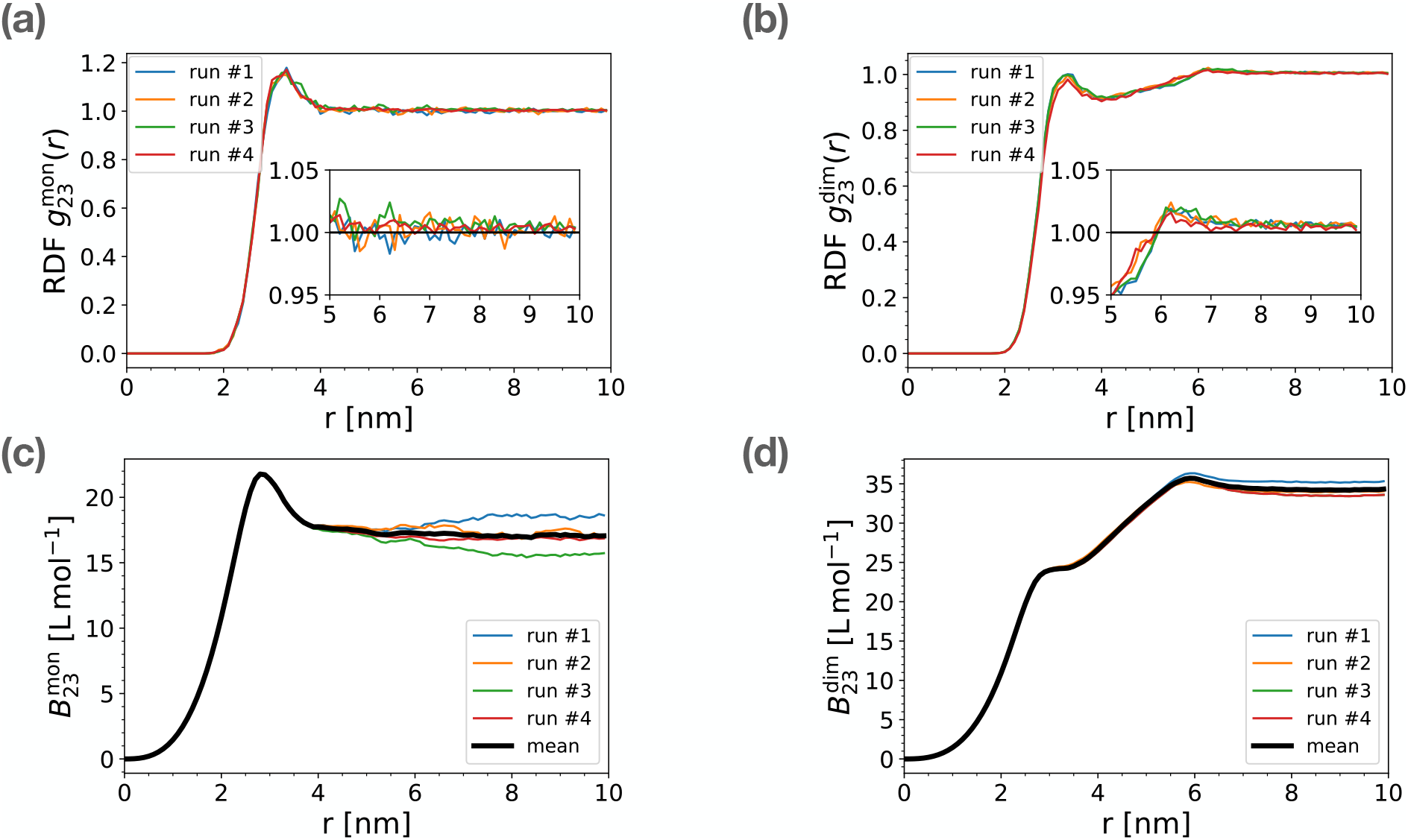
Radial distribution function (RDF) of glucose around monomer and dimer *α*-chymotrypsin and convergence of the associated osmotic second virial coefficient (*B*_23_). The RDFs and *B*_23_ were computed using a protein-sugar scaling parameter *λ* = 0.3 for monomer *α*-chymotrypsin (a, c) and dimer *α*-chymotrypsin (b, d). Insets in panels a and b zoom in on the RDF tail in the range of 5 to 10 nm. For each panel, the results of independent simulation runs are included and the thick black curve in panels c and d shows the average of the runs. To compute 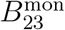 and 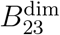, we used Δ = 2 nm and *r*\* = 10 nm.

## References

(1) Sarkar, M.; Li, C.; Pielak, G. J. Soft interactions and crowding. Biophys. Rev. 2013, 5, 187–194.

(2) Zhou, H.-X.; Rivas, G.; Minton, A. P. Macromolecular Crowding and Confinement: Biochemical, Biophysical, and Potential Physiological Consequences. Annu. Rev. Biophys. 2008, 37, 375–397.

(3) Davis-Searles, P. R.; Saunders, A. J.; Erie, D. A.; Winzor, D. J.; Pielak, G. J. Interpreting the effects of small uncharged solutes on protein-folding equilibria. Annu. Rev. Biophys. Biomol. Struct. 2001, 30, 271–306.

(4) Moremen, K. W.; Tiemeyer, M.; Nairn, A. V. Vertebrate protein glycosylation: diversity, synthesis and function. Nature reviews. Molecular cell biology 2012, 13, 448–462.

(5) Langer, M. D.; Guo, H.; Shashikanth, N.; Pierce, J. M.; Leckband, D. E. N-glycosylation alters cadherin-mediated intercellular binding kinetics. J. Cell Sci. 2012, 125, 2478– 2485.

(6) Boushehri, S.; Holey, H.; Brosz, M.; Gumbsch, P.; Pastewka, L.; Aponte-Santamaría, C.; Gräter, F. O-glycans Expand Lubricin and Attenuate its Viscosity and Shear Thinning. Biomacromolecules 2024, 25, 3893–3908.

(7) Lv, P.; Du, Y.; He, C.; Peng, L.; Zhou, X.; Wan, Y.; Zeng, M.; Zhou, W.; Zou, P.; Li, C., et al. O-GlcNAcylation modulates liquid–liquid phase separation of SynGAP/PSD-95. Nat. Chem. 2022, 14, 831–840.

(8) Labokha, A. A.; Gradmann, S.; Frey, S.; Hülsmann, B. B.; Urlaub, H.; Baldus, M.; Görlich, D. Systematic analysis of barrier-forming FG hydrogels from Xenopus nuclear pore complexes. EMBO J. 2013, 32, 204–218.

(9) Vuorio, J.; Škerlová, J.; Fábry, M.; Veverka, V.; Vattulainen, I.; Rězáčová, P.; Martinez-Seara, H. N-Glycosylation can selectively block or foster different receptor-ligand binding modes. Sci. Rep. 2021, 11, 5239, Publisher: Nature Publishing Group.

(10) Saporiti, S.; Laurenzi, T.; Guerrini, U.; Coppa, C.; Palinsky, W.; Benigno, G.; Palazzolo, L.; Ben Mariem, O.; Montavoci, L.; Rossi, M.; Centola, F.; Eberini, I. Effect of Fc core fucosylation and light chain isotype on IgG1 flexibility. Commun. Biol. 2023, 6, 1–10, Publisher: Nature Publishing Group.

(11) Yang, M.; Huang, J.; Simon, R.; Wang, L.-X.; MacKerell, A. D. Conformational Heterogeneity of the HIV Envelope Glycan Shield. Sci. Rep. 2017, 7, 4435, Publisher: Nature Publishing Group.

(12) Marrink, S. J.; Risselada, H. J.; Yefimov, S.; Tieleman, D. P.; De Vries, A. H. The MARTINI force field: coarse grained model for biomolecular simulations. J. Phys. Chem. B 2007, 111, 7812–7824.

(13) Monticelli, L.; Kandasamy, S. K.; Periole, X.; Larson, R. G.; Tieleman, D. P.; Marrink, S.-J. The MARTINI coarse-grained force field: extension to proteins. J. Chem. Theory and Comput. 2008, 4, 819–834.

(14) Marrink, S. J.; Monticelli, L.; Melo, M. N.; Alessandri, R.; Tieleman, D. P.; Souza, P. C. Two decades of Martini: Better beads, broader scope. WIREs Comput. Mol. Sci. 2023, 13, e1620.

(15) Souza, P. C.; Alessandri, R.; Barnoud, J.; Thallmair, S.; Faustino, I.; Grünewald, F.; Patmanidis, I.; Abdizadeh, H.; Bruininks, B. M.; Wassenaar, T. A., et al. Martini 3: a general purpose force field for coarse-grained molecular dynamics. Nat. Meth. 2021, 18, 382–388.

(16) Stark, A. C.; Andrews, C. T.; Elcock, A. H. Toward optimized potential functions for protein–protein interactions in aqueous solutions: osmotic second virial coefficient calculations using the martini coarse-grained force field. J. Chem. Theory Comput. 2013, 9, 4176–4185.

(17) Benayad, Z.; von Bülow, S.; Stelzl, L.; Hummer, G. Simulation of FUS protein condensates with an adapted coarse-grained model. J. Chem. Theory Comput. 2020, 17, 525–537.

(18) Thomasen, F. E.; Pesce, F.; Roesgaard, M. A.; Tesei, G.; Lindorff-Larsen, K. Improving Martini 3 for disordered and multidomain proteins. J. Chem. Theory Comput. 2022, 18, 2033–2041.

(19) Schmalhorst, P. S.; Deluweit, F.; Scherrers, R.; Heisenberg, C.-P.; Sikora, M. Overcoming the limitations of the MARTINI force field in simulations of polysaccharides. J. Chem. Theory Comput. 2017, 13, 5039–5053.

(20) McMillan Jr, W. G.; Mayer, J. E. The statistical thermodynamics of multicomponent systems. J. Chem. Phys. 1945, 13, 276–305.

(21) Cortes-Huerto, R.; Kremer, K.; Potestio, R. Communication: Kirkwood-Buff integrals in the thermodynamic limit from small-sized molecular dynamics simulations. J. Chem. Phys. 2016, 145, 141103.

(22) Köfinger, J.; Hummer, G. Atomic-resolution structural information from scattering experiments on macromolecules in solution. Phys. Rev. E 2013, 87, 052712.

(23) Weatherly, G. T.; Pielak, G. J. Second virial coefficients as a measure of protein-osmolyte interactions. Protein Sci. 2001, 10, 12–16.

(24) Patel, C. N.; Noble, S. M.; Weatherly, G. T.; Tripathy, A.; Winzor, D. J.; Pielak, G. J. Effects of molecular crowding by saccharides on α-chymotrypsin dimerization. Protein Sci. 2002, 11, 997–1003.

(25) Assfalg, M.; Bertini, I.; Del Conte, R.; Giachetti, A.; Turano, P. Cytochrome c and organic molecules: solution structure of the p-aminophenol adduct. Biochemistry 2007, 46, 6232–6238.

(26) Tsukada, H.; Blow, D. Structure of α-chymotrypsin refined at 1.68 Å resolution. J. Mol. Biol. 1985, 184, 703–711.

(27) de Jong, D. H.; Singh, G.; Bennett, W. D.; Arnarez, C.; Wassenaar, T. A.; Schafer, L. V.; Periole, X.; Tieleman, D. P.; Marrink, S. J. Improved parameters for the martini coarse-grained protein force field. J. Chem. Theory Comput. 2013, 9, 687–697.

(28) Kabsch, W.; Sander, C. Dictionary of protein secondary structure: pattern recognition of hydrogen-bonded and geometrical features. Biopolymers 1983, 22, 2577–2637.

(29) Periole, X.; Cavalli, M.; Marrink, S.-J.; Ceruso, M. A. Combining an elastic network with a coarse-grained molecular force field: structure, dynamics, and intermolecular recognition. J. Chem. Theory Comput. 2009, 5, 2531–2543.

(30) Abraham, M. J.; Murtola, T.; Schulz, R.; Páll, S.; Smith, J. C.; Hess, B.; Lindahl, E. GROMACS: High performance molecular simulations through multi-level parallelism from laptops to supercomputers. SoftwareX 2015, 1, 19–25.

(31) Humphrey, W.; Dalke, A.; Schulten, K. VMD: visual molecular dynamics. J. Mol. Graph. 1996, 14, 33–38.

(32) De Jong, D. H.; Liguori, N.; Van Den Berg, T.; Arnarez, C.; Periole, X.; Marrink, S. J. Atomistic and coarse grain topologies for the cofactors associated with the photosystem II core complex. J. Phys. Chem. B 2015, 119, 7791–7803.

(33) Dolinsky, T. J.; Czodrowski, P.; Li, H.; Nielsen, J. E.; Jensen, J. H.; Klebe, G.; Baker, N. A. PDB2PQR: expanding and upgrading automated preparation of biomolecular structures for molecular simulations. Nucleic Acids Res. 2007, 35, W522–W525.

(34) Jurrus, E.; Engel, D.; Star, K.; Monson, K.; Brandi, J.; Felberg, L. E.; Brookes, D. H.; Wilson, L.; Chen, J.; Liles, K., et al. Improvements to the APBS biomolecular solvation software suite. Protein Sci. 2018, 27, 112–128.

(35) Wang, J.; Cieplak, P.; Kollman, P. A. How well does a restrained electrostatic potential (RESP) model perform in calculating conformational energies of organic and biological molecules? J. Comput. Chem. 2000, 21, 1049–1074.

(36) Jo, S.; Kim, T.; Iyer, V. G.; Im, W. CHARMM-GUI: a web-based graphical user interface for CHARMM. J. Comput. Chem. 2008, 29, 1859–1865.

(37) Lee, J.; Hitzenberger, M.; Rieger, M.; Kern, N. R.; Zacharias, M.; Im, W. CHARMM-GUI supports the Amber force fields. J. Chem. Phys. 2020, 153 .

(38) Bussi, G.; Donadio, D.; Parrinello, M. Canonical sampling through velocity rescaling. J. Chem. Phys. 2007, 126, 014101.

(39) Berendsen, H. J.; Postma, J. v.; van Gunsteren, W. F.; DiNola, A.; Haak, J. R. Molecular dynamics with coupling to an external bath. J. Chem. Phys. 1984, 81, 3684–3690.

(40) Parrinello, M.; Rahman, A. Polymorphic transitions in single crystals: A new molecular dynamics method. J. Appl. Phys. 1981, 52, 7182–7190.

(41) Mugnai, M. L.; Shin, S.; Thirumalai, D. Entropic contribution of ACE2 glycans to RBD binding. Biophys. J. 2023,

(42) von Bülow, S.; Sikora, M.; Blanc, F. E.; Covino, R.; Hummer, G. Antibody accessibility determines location of spike surface mutations in SARS-CoV-2 variants. PLoS Comput. Biol. 2023, 19, e1010822.

(43) Luthey-Schulten, Z. Integrating experiments, theory and simulations into whole-cell models. Nat. Meth. 2021, 18, 446–447.

## References

(S1) Dolinsky, T. J.; Czodrowski, P.; Li, H.; Nielsen, J. E.; Jensen, J. H.; Klebe, G.; Baker, N. A. PDB2PQR: expanding and upgrading automated preparation of biomolecular structures for molecular simulations. Nucleic Acids Res. 2007, 35, W522–W525.

(S2) Jurrus, E.; Engel, D.; Star, K.; Monson, K.; Brandi, J.; Felberg, L. E.; Brookes, D. H.; Wilson, L.; Chen, J.; Liles, K., et al. Improvements to the APBS biomolecular solvation software suite. Protein Sci. 2018, 27, 112–128.

(S3) Wang, J.; Cieplak, P.; Kollman, P. A. How well does a restrained electrostatic potential (RESP) model perform in calculating conformational energies of organic and biological molecules? J. Comput. Chem. 2000, 21, 1049–1074.

(S4) Jo, S.; Kim, T.; Iyer, V. G.; Im, W. CHARMM-GUI: a web-based graphical user interface for CHARMM. J. Comput. Chem. 2008, 29, 1859–1865.

(S5) Lee, J.; Hitzenberger, M.; Rieger, M.; Kern, N. R.; Zacharias, M.; Im, W. CHARMM-GUI supports the Amber force fields. J. Chem. Phys. 2020, 153 .

(S6) Schmalhorst, P. S.; Deluweit, F.; Scherrers, R.; Heisenberg, C.-P.; Sikora, M. Overcoming the limitations of the MARTINI force field in simulations of polysaccharides. J. Chem. Theory Comput. 2017, 13, 5039–5053.

(S7) Benayad, Z.; von Bülow, S.; Stelzl, L.; Hummer, G. Simulation of FUS protein condensates with an adapted coarse-grained model. J. Chem. Theory Comput. 2020, 17, 525–537.

